# Anti-consensus: detecting trees that have an evolutionary signal that is lost in consensus

**DOI:** 10.1101/706416

**Authors:** Daniel H. Huson, Benjamin Albrecht, Sascha Patz, Mike Steel

## Abstract

In phylogenetics, a set of gene trees is often summarized by a consensus tree, such as the majority consensus, which is based on the set of all splits that are present in more than 50% of the input trees. A “consensus network” is obtained by lowering the threshold and considering all splits that are contained in 10% of the trees, say, and then computing the corresponding splits network. By construction and in practice, a consensus network usually shows the majority tree, extended by a number of rectangles that represent local rearrangements around internal nodes of the consensus tree. This may lead to the false conclusion that the input trees do not differ in a significant way because “even a phylogenetic network” does not display any large discrepancies. To harness the full potential of a phylogenetic network, we introduce the new concept of an anti-consensus network that aims at representing the largest interesting discrepancies found in a set of gene trees. We provide an efficient algorithm for computing an anti-consensus and illustrate its application using a set of gene trees from species and strains in the genus *Kosakonia*.

## Introduction

The main aim of phylogenetic analysis is to determine the *species tree T*_*species*_ that describes the evolutionary history of a set of organisms. One approach to this is to first estimate a set of *genes trees T*_1_, …, *T*_*m*_ for the organisms and then to obtain the species tree as a consensus tree of the gene trees (Margush and McMorris, 1981).

This type of majority tree *T*_*maj*_ is obtained from the set 𝒮_*h*_ of all “splits” that occur in more than *h* = 50% of the gene trees, which is always compatible and can be represented by a uniquely defined phylogenetic tree, namely the majority tree (Bryant, 2003). If the threshold *h* is set below 50%, then the resulting set of splits 𝒮_*h*_ need not be compatible, in which case it will not be representable by a phylogenetic tree but instead by a more general splits network (Bandelt and Dress, 1992). A network obtained in this way is called a *consensus network* and was introduced by Holland et al. (2004).

At first glance, the concept of a consensus network seems promising, as one might hope that such a network will display any interesting patterns of discordance that exist among the input trees. However, in practice, consensus networks usually provide very little additional insight: they emphasize local disagreements among the input trees that are shared by a substantial proportion of the trees while suppressing major discrepancies that arise in small numbers of trees. Indeed, the example presented by Holland et al. (2004) has only two minor rearrangements around two different internal nodes. See also Figures 1(f) and 2 below.

**Fig. 1.**
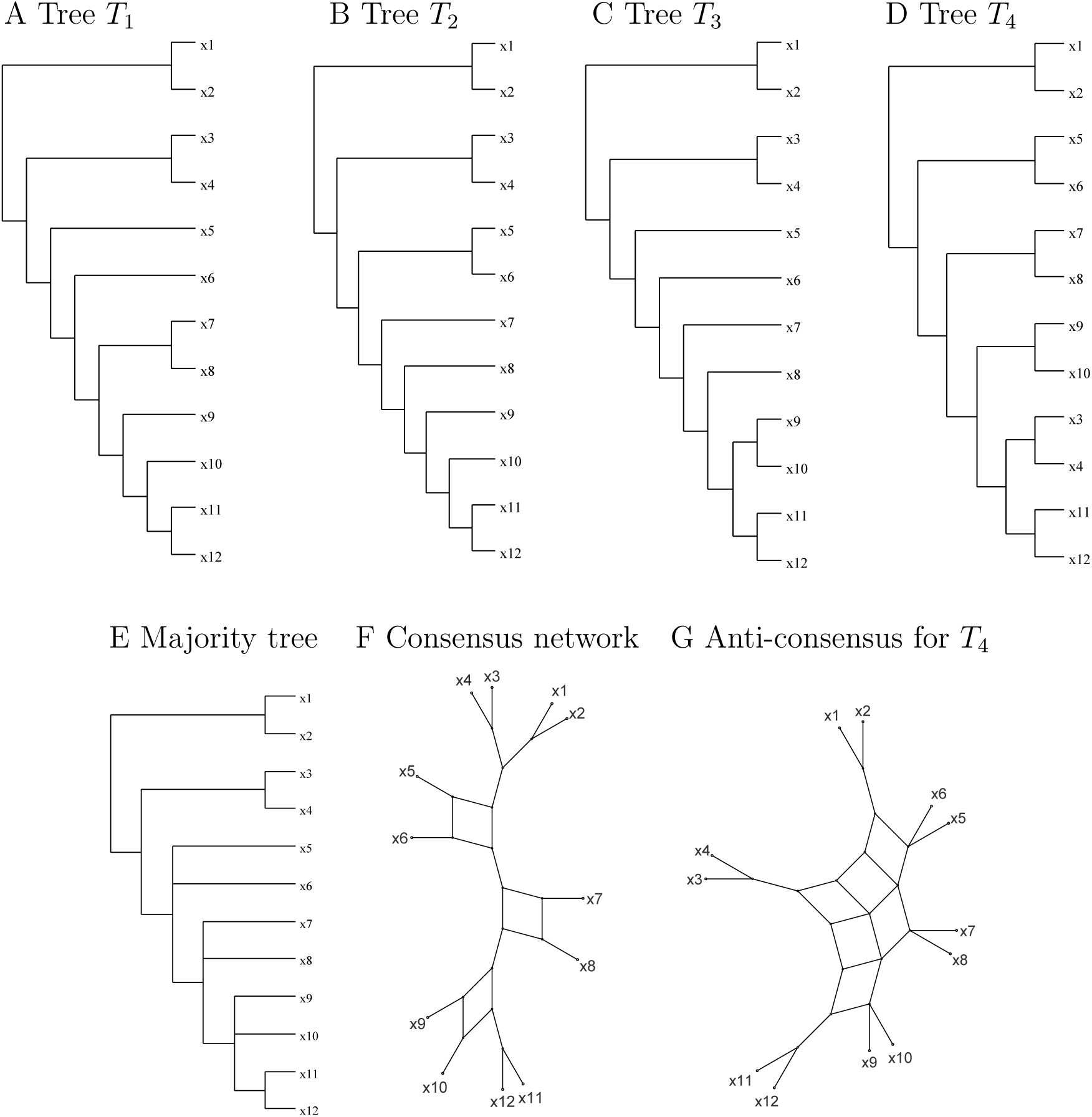
For four hypothetical input trees *T*_1_, …, *T*_4_ displayed in (A–D), we show (E) the majority tree *T*_*ref*_, (F) the consensus network using a 30% threshold, and (G) the unique (single-event) anti-consensus network for this data, which is associated with tree *T*_4_.

Here, we introduce the new concept of an *anti-consensus* network that aims at representing the “interesting” large discrepancies present in a collection of gene trees while suppressing the “uninteresting” discrepancies that are caused by lack of resolution (e.g. short branches, which lead to local rearrangements) or random noise. We are particularly interested in detecting discrepancies that may be caused by large horizontal gene transfer events.

Determining the minimum number of horizontal gene transfer events required to reconcile two phylogenetic trees is computationally hard (Humphries et al., 2013). Here, we describe an anti-consensus method that is based on the concept of distortion (Huson et al., 2006) and heuristically seeks to detect large discrepancies between an individual input tree and a given reference tree, such as the majority consensus of all input trees or, say, an established species tree. We demonstrate the application of the method on a collection of 2 325 gene trees on 29 different species and strains of the bacterial genus *Kosakonia*, and explore its performance in a simulation study. The anti-consensus method is implemented in our new program SplitsTree5, which is currently under development and available from www.splitstree.org.

## Material and Methods

### Preliminaries

In the following, let *X* be a set of *n* taxa. Recall that a split *S* = *A* | *B* on *X* is a bipartition of *X* into two non-empty and disjoint sets *A* and *B* whose union equals *X* (Bandelt and Dress, 1992). Let *T* be a phylogenetic tree on the leaf set *X*. Deletion of any edge *e* in *T* separates the set of taxa *X* into a split *S*_*e*_ = *A* | *B* that is uniquely associated with *e*. The set of all such splits (each associated with one edge of *T)* is called the *split encoding* of *T*, denoted 𝒮(*T)*. Two splits *S* and *S*′ on *X* are called *compatible*, if at least one of the four possible intersections of either part of *S* with either part of *S*′ is empty. This condition is equivalent to requiring that both splits appear in the split encoding of the same phylogenetic tree (Buneman, 1974). We say that a split *S* is *compatible* with a tree *T* on *X* if *S* is compatible with all the splits in the split encoding 𝒮(*T)* of *T*. This condition holds if and only if *S* is contained in 𝒮(*T)* or if a refinement *T* ′ of *T* exists such that *S* is contained in 𝒮(*T* ′) (Buneman, 1974).

Let *T*_1_, …, *T*_*m*_ be a set of *m* phylogenetic trees on *X*. We will assume that all input edges have weights and that the trees are normalized (i.e., that the total weight of all edges in any input tree is 1). Additionally, we require a *reference* tree *T*_*ref*_ on the same set of taxa. This might be the *majority tree* of the given input trees, or a “species tree” established in some way.

We use 𝒮_*ref*_ to denote the set of splits associated with the reference tree. If we are using the majority consensus tree as the reference tree, then 𝒮_*ref*_ will contain all of the splits that are associated with more than 50% of the input trees.

Let *S* and *S*′ be two input splits that are both incompatible with the reference tree. We say that *S covers S*′ (with respect to *T*_*ref*_) if *S* is incompatible with all splits in 𝒮_*ref*_ with which *S*′ is also incompatible. Note that it can happen that two different splits *S* and *S*′ are both incompatible with exactly the same set of splits in 𝒮_*ref*_. In this case, for *S* to cover *S*′, we also require *S* to be lexicographically smaller than *S*′. We use *C*(*S*) to denote the set of all such splits *S*′ that are covered by *S*, together with *S*.

The *distortion* of a split *S* with respect to a phylogenetic tree *T*, denoted by *D*(*S, T)*, is the minimum number of *subtree prune and re-graft* operations that must be applied to *T* to obtain a phylogenetic tree that is compatible with *S*. We introduced this concept in Huson et al. (2006) and showed how to compute this number efficiently (in time polynomial in *n* = |*X*|). Note that a split *S* is compatible with a tree *T* if and only if *D*(*S, T)* = 0.

### The anti-consensus

We now introduce the concepts required for defining a first ‘anti-consensus’ method. Let *T*_*i*_ be one of the input trees. We would like to identify a set of splits in *T*_*i*_ that would be expected to arise in a large horizontal gene transfer event involving the gene on which *T*_*i*_ is based. Such a set of <u>s</u>ignificantly <u>in</u>compatible splits will be called a *SIN*. To define a SIN more precisely, let *D*_1_(*T*_*i*_, *T*_*ref*_) denote the set of all splits in *T*_*i*_ that have a distortion value of 1 with respect to the reference tree. Given a split *S* in *D*_1_(*T*_*i*_, *T*_*ref*_) that is *not covered* by any other split in *D*_1_(*T*_*i*_, *T*_*ref*_), a SIN *Z* is obtained by taking split *S* along with all the other splits in *D*_1_(*T*_*i*_, *T*_*ref*_) that are covered by *S*. In other words, for an input tree *T*_*i*_, we have:

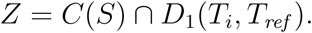

We now discuss how to score and compare different SINs. The *incompatibility score γ*(*Z*) of a SIN *Z* is defined as the sum of the weights of all splits in the reference tree that are incompatible with any split in *Z*. It follows from the definition of a SIN that it suffices to consider the one uncovered split contained in *Z* to calculate this. Further, the *weight ω*(*Z*) of a SIN *Z* is given by the sum of the weights of all splits in *Z*, namely:

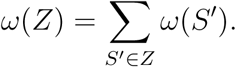

The *(single-event) anti-consensus* for a set of gene trees *T*_1_, …, *T*_*m*_ on *X* is defined as the list of all associated SINs *Z*_1_, *Z*_2_, …, ranked by decreasing incompatibility score. In practice, it makes sense to use a threshold to ignore SINs with a very low weight.

The computation of the single-event anti-consensus is based on a polynomial number of pair-wise split compatibility determinations and distortion calculations, both of which are operations that can be performed in polynomial time. Hence, the calculation of all SINs can be completed in polynomial time.

Each SIN *Z* can be represented by the splits network computed for the union of *Z* and the set of reference splits 𝒮_*ref*_ (see Figure 1G).

### Conceptual example

To illustrate the newly introduced concepts, we first consider a hypothetical example consisting of four input trees *T*_1_, …, *T*_4_ on taxa *X* = {*x*_1_, …, *x*_12_}, as shown in Figure 1A–D). We will use the majority consensus tree as the reference tree.

The majority tree for these four taxa is shown in Figure 1E and the consensus network based on a threshold of 30% is shown in Figure 1F. The consensus network indicates (only) that there are local rearrangements around three internal nodes associated with the pairs of taxa *x*_5_ and *x*_6_, *x*_7_ and *x*_8_, and *x*_9_ and *x*_10_, respectively.

In this made-up example, tree *T*_4_ differs from the other three trees by the positioning of the taxon pair (*x*_3_, *x*_4_). In trees *T*_1_, *T*_2_ and *T*_3_, this taxon pair is adjacent to the pair (*x*_1_, *x*_2_), whereas in tree *T*_4_, the pair is positioned adjacent to the pair (*x*_11_, *x*_12_). As a result, in the reference tree *T*_*ref*_, the pair (*x*_3_, *x*_4_) appears adjacent to (*x*_1_, *x*_2_). This conflicting placement of the pair (*x*_3_, *x*_4_) in *T*_*ref*_ and *T*_4_ gives rise to three splits that belong to *D*_1_(*T*_4_, *T*_*ref*_), namely *S*_1_ = {*x*_1_, *x*_2_, *x*_5_, *x*_6_} vs. the rest, *S*_2_ = {*x*_1_, *x*_2_, *x*_5_, *x*_6_, *x*_7_, *x*_8_} vs. the rest, and *S*_3_ = {*x*_1_, *x*_2_, *x*_5_, *x*_6_, *x*_7_, *x*_8_, *x*_9_, *x*_10_} vs. the rest.

The split *S*_1_ is incompatible with the three main internal splits down the backbone of the reference tree, whereas *S*_2_ and *S*_3_ are incompatible with two and one of them, respectively. The set *Z* = {*S*_1_, *S*_2_, *S*_3_} is a SIN associated with tree *T*_4_. It is incompatible with three of the six internal splits of *T*_*ref*_, so its incompatibility score is 50%.

The (single-event) anti-consensus network shown in Figure 1G is the splits network representing the union of the set of splits in *T*_*ref*_ and *Z*. It clearly indicates the ambiguous placement of the pair of taxa (*x*_3_, *x*_4_). In contrast, the consensus network shown in Figure 1F gives no indication that a large discrepancy among the four input trees exists.

### *Application to the bacterial genus* Kosakonia

*Kosakonia* is a novel genus that contains a number of species that were originally assigned to the genus *Enterobacter* (Li et al., 2016). It is of particular interest because it contains plant growth-promoting strains as well as potential human pathogens.

We used PanX (Ding et al., 2018) (using the default mode recommended for small datasets) to identify 2 325 genes that are all shared by 29 species and strains of the genus *Kosakonia*. For each gene, we downloaded the 29 sequences from PanX and then aligned them with MUSCLE (Edgar, 2004) (v3.8.31, default settings, -maxiters 8). The alignments were trimmed with trimAl (Capella-Gutierrez et al., 2009) (v1.2, parameters: -gt 0.8 –st 0.001 -cons 60). Gene trees were inferred with FastTree (Price et al., 2009) (v.2.1, parameters: -gtr -nt). In addition, we downloaded the *Kosakonia* “strain tree” provided by PanX. All trees are available here:http://ab.inf.uni-tuebingen.de/data/software/splitstree5/download/anti-consensus-trees.zip

First, we computed the consensus network for all 2 325 trees with SplitsTree5, keeping all splits that appear in more than 5% of all input trees. This is displayed in Figure 2. This is an example of a consensus network that emphasizes local rearrangements around central nodes.

**Fig. 2.**
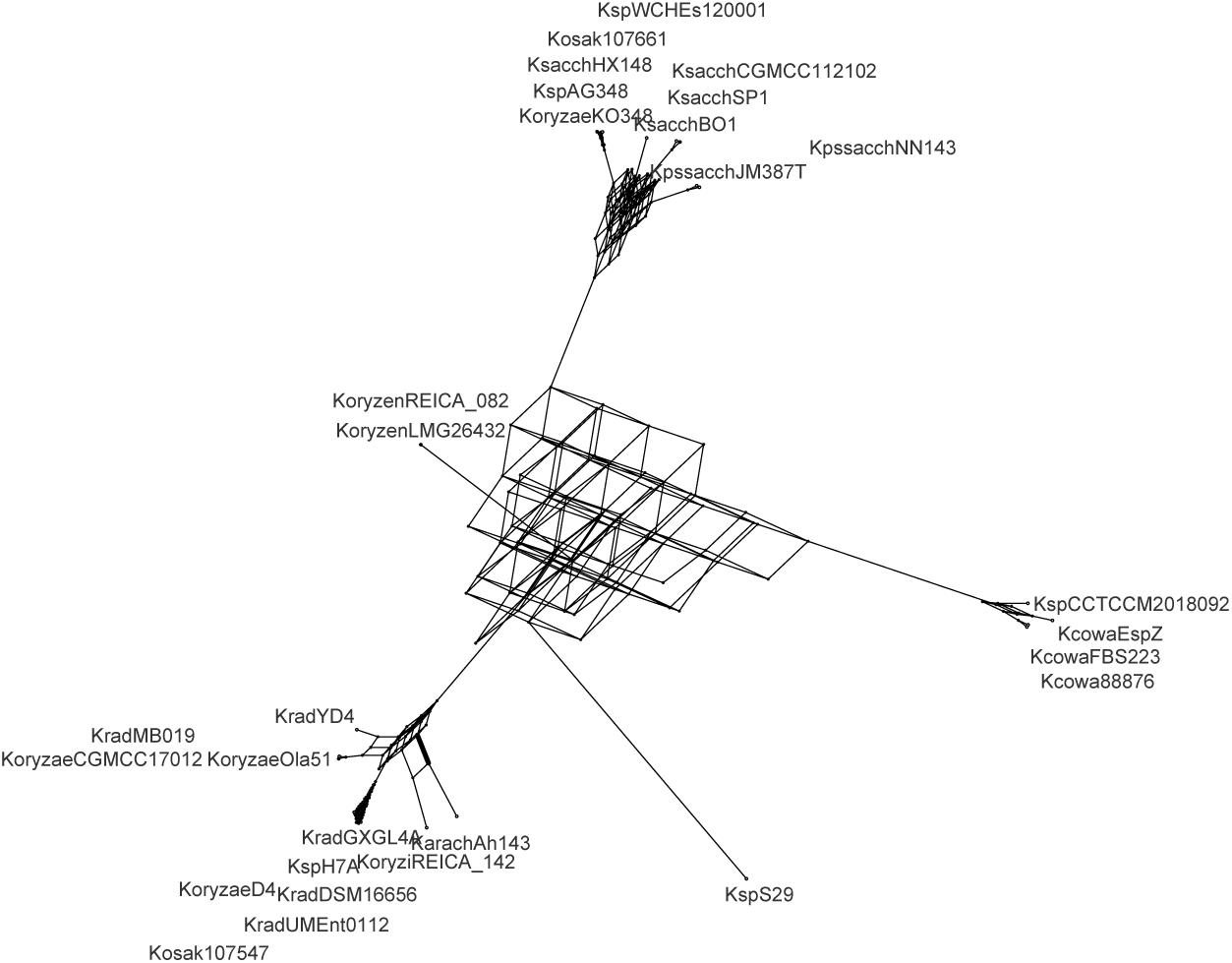
A consensus network for 2 325 gene trees on the genus of *Kosakonia*. This represents all splits that occur in more than 5% of all input trees.

We ran the single-event anti-consensus method on the set of 2 325 trees, using default parameters and using the downloaded strain tree as reference tree. This took less than 1 s on a laptop with 12 cores.

There are 60 SINs whose incompatibility scores are ≥ 10% of the total weight of all internal edges of the reference tree. We list the top 10 SINs in Table 1. We present the anti-consensus networks associated with these ten SINs in Figure 3. In each case, the networks show that the discrepancy between the corresponding gene tree and the reference tree spans a large proportion of the reference tree cleary.

**Table 1.** Top 10 anti-consensus SINs from 2 325 genes from 29 *Kosakonia* strains. For each SIN, we report (a) its rank, (b) the incompatibility score as a percentage of the total length of the reference tree, (c) the total weight and (d) the number of splits in the SIN, (e) the tree number in file, (f) the gene name and (g) the associated protein.

| (a)<br>SIN | (b)<br>Score | (c)<br>Weight | (d)<br>Splits | (e)<br>Tree | (f)<br>Gene | (g)<br>Protein |
| --- | --- | --- | --- | --- | --- | --- |
| 1 | 22.5% | 0.4363 | 3 | 2312 | rplN | LSU ribosomal Protein L14p |
| 2 | 21.3% | 0.1657 | 5 | 1055 | impD | HCP1 family type VI secretion system effector protein |
| 3 | 21.0% | 0.0217 | 2 | 1481 | rpsO | SSU ribosomal protein S15p |
| 4 | 19.4% | 0.1664 | 4 | 2322 | rpsJ | SSU ribosomal protein S10p |
| 5 | 18.1% | 0.3268 | 5 | 2317 | rplV | LSU ribosomal protein L22p |
| 6 | 18.0% | 0.0640 | 1 | 2314 | rpmC | LSU ribosomal protein L29p |
| 7 | 17.5% | 0.0753 | 1 | 1459 | rpsU | SSU ribosomal protein S21p |
| 8 | 17.5% | 0.0063 | 1 | 410 | ompF | Outer membrane protein F |
| 9 | 16.8% | 0.1005 | 3 | 1891 | rpsP | SSU ribosomal protein S16p |
| 10 | 16.8% | 0.0936 | 3 | 2313 | rpsQ | SSU ribosomal protein S17p |

**Fig. 3.**
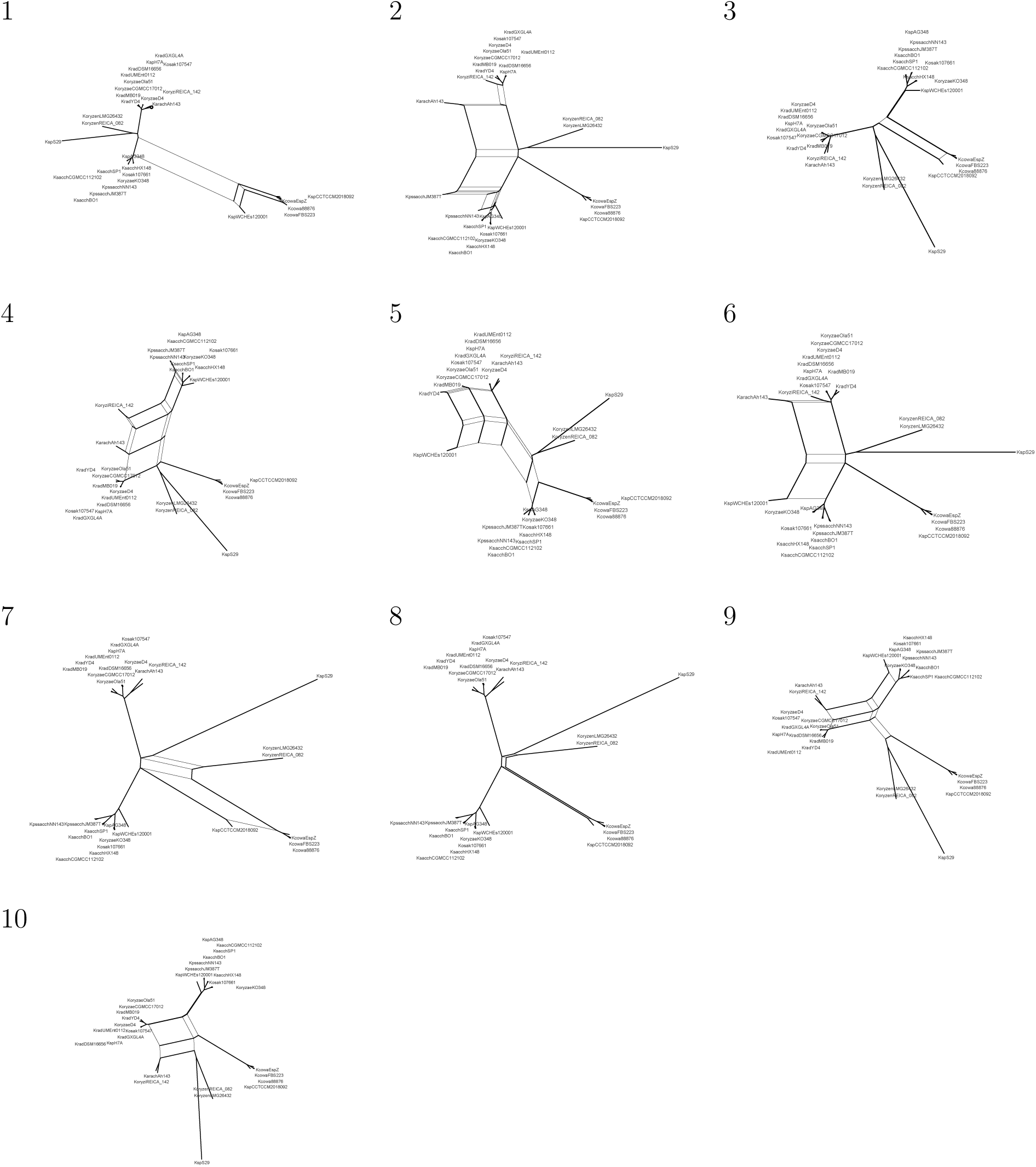
Anti-consensus networks. The ten anti-consensus networks corresponding to the top 10 SINs reported in the main Table.

Eight of the ten listed SINs correspond to ribosomal proteins. One exception is SIN 2, which corresponds to a HCP1 family type VI secretion system effector protein, considered a virulence factor in some pathogens. The other exception is SIN 8, which corresponds to the outer membrane protein F, which forms a pore and has been associated with antibacterial mechanisms. For these two genes of particular interest, we provide a tanglegram (Huson and Scornavacca, 2012) of the gene tree and the reference tree to clearly show the large difference between the gene trees and reference tree, see Figure 4.

**Fig. 4.**
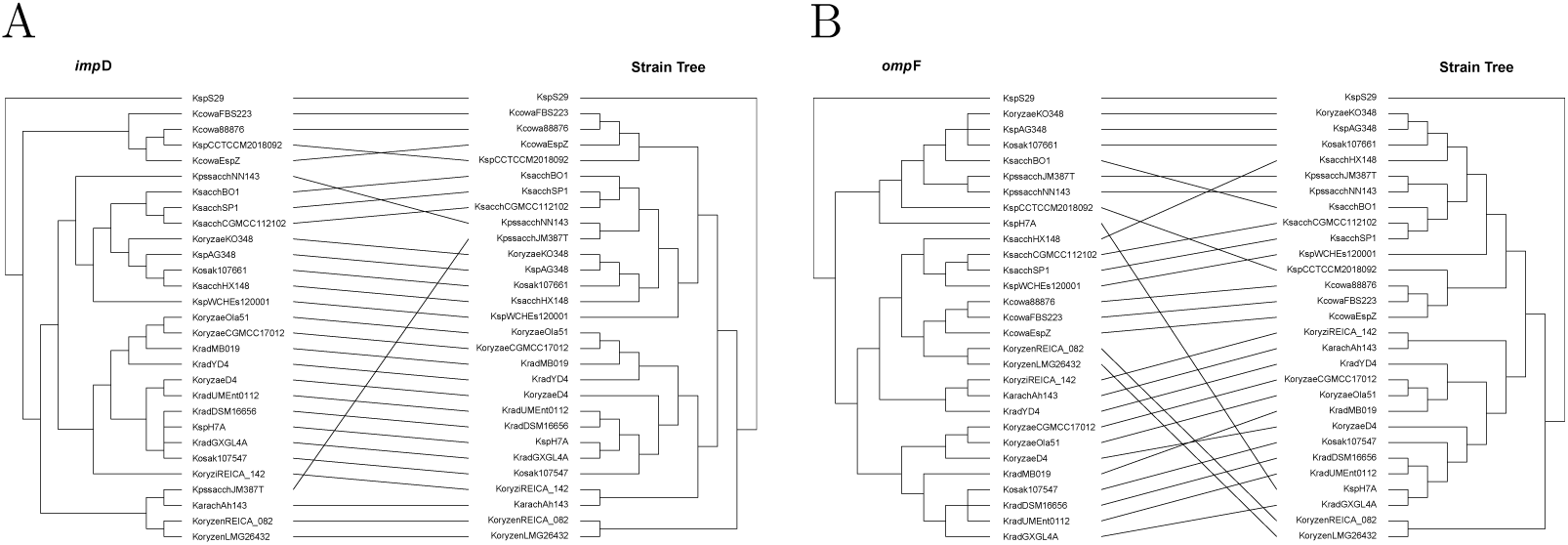
Tanglegrams. For SIN 2 and SIN 8 listed in the main Table, corresponding to the genes ImpD and ompF, we display a tanglegram of the corresponding gene tree and the reference tree, in A and B, respectively.

### Simulation study

How well does the anti-consensus identify large HGT events in a collection of incongruent gene trees? To investigate this question in a simulation study, we used the program SimPhy (Mallo et al., 2016) to generate many different profile of trees that differ topologically as result of incomplete lineage sorting, with the amount of difference determined by the program’s “population size” parameter. A subset of the trees of each such simulated tree profile was then modified by applying large *rooted subtree-prune-and-regraft* (rSPR) operations to each of them, so as to obtain a set of *target trees* that harbor the HGT events to be found.

In more detail, we performed a single-event simulation as follows. For a fixed population size ranging from 10 to 250, 000, we used SimPhy to create a set of 1, 000 different gene trees on a fixed set of 250 taxa. The first 10 gene trees were then used as the set of *target trees* and each was subjected to a single “long-range” rSPR modification that covered at least 70% of the total diameter of the tree. This construction was repeated 100 times for each choice of population size.

The single-event method was then applied to each of these simulated datasets. For each result, a target tree gave rise to a *true positive*, if it occurred within the top 10 of all computed SINs. Based on this, we report the average true positive rates (TP) as a function of population size in Figure 5. Note that in this simple setup, both the false positive rate and the false negative rate are equal to 1 − TP.

**Fig. 5.**
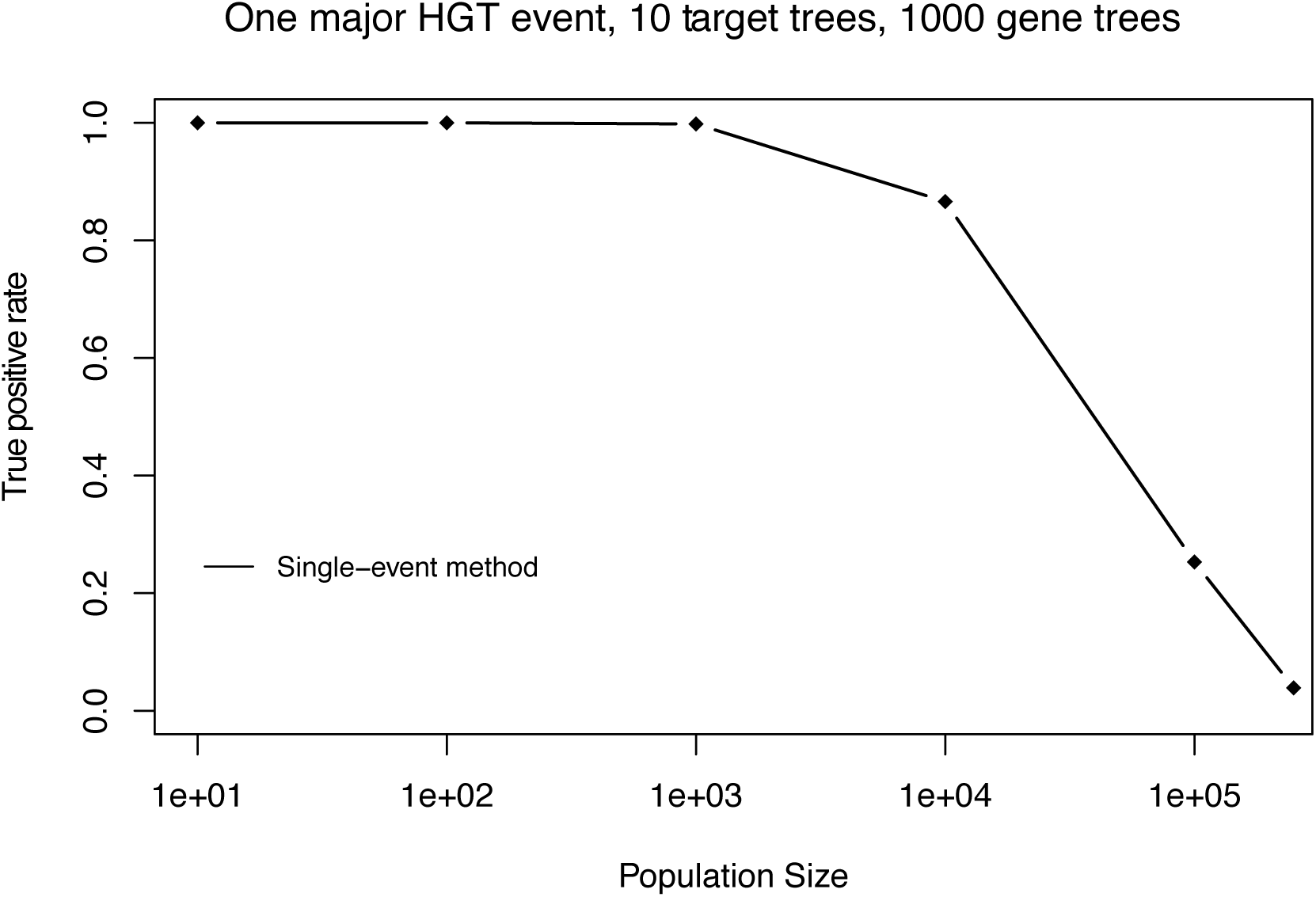
Simulation study of the single-event method. Here we present the true positive rate for the single-event anti-consensus as a function of SimPhy’s “population size” parameter (100 replicates per value). We simulated 1000 trees on 250 taxa under a model of incomplete lineage sorting and then modified 10 target trees so as to each include a large HGT event. A target tree is deemed a true positive, if and only if it mentioned among the first ten reported SINs.

### Multiple-event anti-consensus

The single-event anti-consensus heuristic described above addresses the situation in which one or more of the input trees involves a gene that has been affected by a single large HGT event (with different HGT events on different trees indicated by different SINs). In practice, a single input tree might be affected by many such events.

If two such events affect two different parts of the involved gene tree *T*_*i*_ and two different parts of the reference tree *T*_*ref*_, respectively, then the heuristic described above will, in fact, detect the two corresponding disjoint SINs.

If, however, two or more HGT events interfere with each other, then this will give rise to splits with a distortion value of ≥ 2, with respect to *T*_*ref*_. In this case, the heuristic described above might not detect a corresponding SIN.

A simplistic approach to dealing with multiple HGT events that affect the same part of an input tree *T*_*i*_ is to search for a split *S* in *T*_*i*_ that has a distortion value of ≥ 1 with respect to *T*_*ref*_ and is not covered by any other split in *T*_*i*_. Thus, a *(multi-event) SIN Z* is given by taking *S*, together will all splits covered by *S*; in other words, using *Z* = *C*(*S*).

In practice, because there is no restriction on the distortion value of the splits considered here, this approach will be more sensitive, as it is able to pick up large HGT events in the presence of noise, but it is less specific, potentially producing high-ranking SINs that are not caused by large HGT events.

In addition, another consequence of having no restriction on the distortion value of the splits is that many similar SINs for the same input tree *T*_*i*_ may occur. To address this issue, we disregard any SIN *Z* for which there is another SIN *Z*′ that has a higher incompatibility score and is incompatible with at least one of the reference splits with which *Z* is incompatible. In addition, in our implementation, we also provide a mode in which at most one SIN is reported per input tree.

The *multi-event* anti-consensus method determines all (multi-event) SINs in polynomial time, and any such SIN *Z* can be represented by the splits network computed for the union of *Z* and the set of reference splits 𝒮_*ref*_.

*Multiple-event simulation* We performed a multi-event simulation to evaluate the performance of our approach in the presence of multiple, interfering HGT events on a single input tree. Here, we repeated the simulation procedure described above, except that we applied five long-range rSPR modifications to each of the 10 target trees. We report the average true positive rates as a function of population size in Figure 6.

**Fig. 6.**
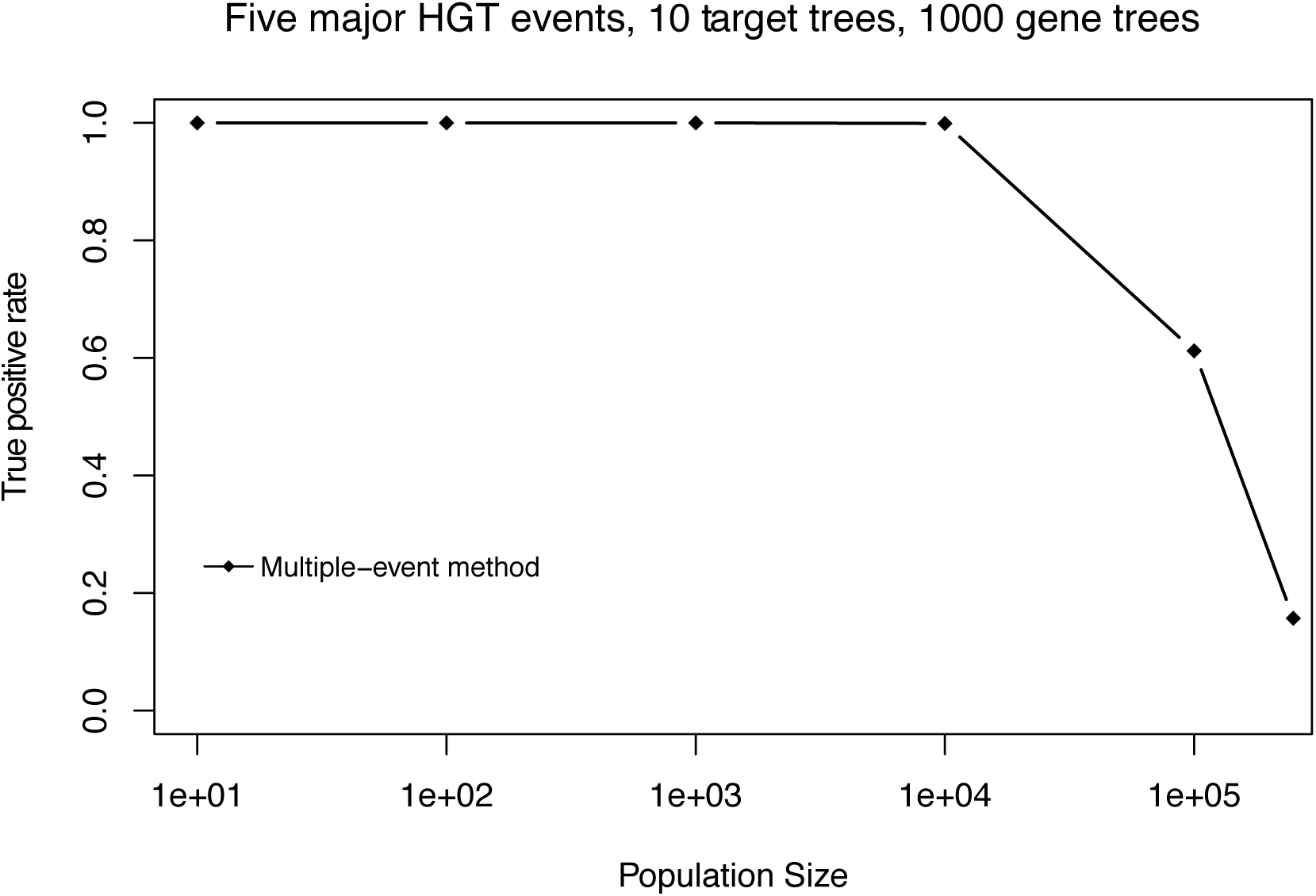
Simulation study of the multi-event method. Here, we present the true positive rate for the multiple-event anti-consensus method as a function of SimPhy’s “population size” parameter (100 replicates per value). We simulated 1000 trees on 250 taxa under a model of incomplete lineage sorting and then modified 10 target trees so each included five large horizontal gene transfer events. A target tree is deemed to be a true positive, if and only if it is mentioned among the first 10 reported significantly incompatible splits.

## Results

We have implemented both the single-event anti-consensus and the multiple-event anti-consensus method in our new program SplitsTree5, which is under development and freely available at www.splitstree.org. The algorithm is implemented in parallel and thus required less than 1 s to process 2 325 trees on 29 taxa. We have run the algorithm on other datasets containing thousands of trees and hundreds of taxa, observing running times of only a few seconds.

Application of the algorithm to 2 325 gene trees on 29 strains and species of *Kosakonia* identifies 60 gene trees that differed from the species tree by an incompatibility score of more than 10%. Among the highest scoring SINs, we found two genes that are potentially associated with pathogenecity or antibacterial mechanisms.

The simulation study shows that the ability of the anti-consensus method to detect gene trees that differ because of major HGT events depends on the amount of phylogenetic “noise” in the set of trees, which, in our simulation study, is represented by the population size used by the SimPhy tree simulator. For low levels of noise, the anti-consensus method consistently ranks the SINs associated with the target trees first. As the noise level increases, trees with discrepancies caused by noise compete with the target trees.

## Discussion

In this age of high-throughput sequencing, the phylogenetic analysis of a set of organisms can be based on thousands of genes. A researcher might use a a consensus network method in the hope that the resulting phylogenetic network will reveal large discrepancies caused by HGT. However, by definition, consensus methods aim at suppressing large events that affect only a small number of trees.

Here, we have proposed the new concept of an anti-consensus network that aims at emphasizing large discrepancies in the input data that are of interest, while suppressing smaller differences in the input data, such as local rearrangements around internal nodes of the consensus tree that may result from mechanisms such as incomplete lineage sorting, or may reflect inaccuracies in the inferred trees.

The anti-consensus methods introduced in this paper can be computed efficiently, even on large datasets involving thousands of trees and hundreds of taxa. We applied the method to a collection of bacterial strains and species of the genus *Kosakonia*. To illustrate the kind of output that the method provides, we list the 10 most incompatible genes and show anti-consensus networks for two of the gene trees.

The anti-consensus method may also provide a convenient filter to identify and remove genes trees that differ strongly from a proposed species tree, so as to allow a better estimation of the species tree using only those gene trees that do not appear to have been subjected to large HGT events.

